# Integrating phylodynamics and historical records reveals decades-old introductions of PRRSV into Costa Rica via international swine trade

**DOI:** 10.1101/2025.08.22.671733

**Authors:** Bernal León, Samiah Kanwar, Olga Aguilar, Idania Chacón, Gisela Cháves, David J. Spiro, Sana Tamim, Nidia S. Trovao

## Abstract

**Background:** In August 1995, necropsies on post-weaning piglets from the CA−CART farm in the province of Cartago, Costa Rica, revealed respiratory lesions, pleuritis, peritonitis, and arthritis. Skin lesions were also observed, progressing to scabs. A subsequent outbreak in 1996 prompted antibiotic administration. Mortality analysis from 1990 to 1995 showed no significant seasonal patterns, but yearly variations were noted. Piglets born in Costa Rica from imported gilts had a higher average mortality rate (10.65%) than the 8.11% mortality rate for piglets born from non-imported gilts and sows or imported sows (p = 0.002).

**Methods:** In March 1996, serum samples were sent for potential PRRS virus (PRRSV) diagnosis, and 10 PRRSV-2 ORF5 sequences collected during a prospective study from 2019-2021 were obtained from various locations in the country. In this article, we seek to investigate the evolutionary and spatio-temporal dynamics of PRRSV-2 in Costa Rica in the context of international swine movements using the phylodynamic framework integrated with the BEAST package.

**Results and Discussion:** The phylodynamic modeling estimated at least two independent PRRSV-2 introductions into the country. The earliest introduction occurred around 1978 (95% highest posterior density interval: 1959.01-1997.54) and led to the viruses circulating in the farm 1CRC with an origin from Japan, possibly via US swine exports. A second cluster, 4CRC, subsequently emerged within Costa Rica from the earlier 1CRC lineage (1990.24; 95% HPD interval = 1976.15-2010.30). Another viral introduction from the US occurred around 1991 (95% highest posterior density interval: 1974.34-2011.35) and established the 3CRC cluster. Though the viral introduction was traced back to the US, the limited genomic surveillance of PRRSV leaves room for considering alternative origins.

**Conclusion:** Our findings provide a high-resolution model of how international swine trade drives the introduction and evolution of a major livestock pathogen, highlighting the critical need for integrating genomic surveillance into biosecurity protocols.

## INTRODUCTION

The porcine reproductive and respiratory virus (PRRSV) has had a significant negative economic impact on swine production since the first cases were detected in Iowa pig pens in 1985 [1]. For example, in the United States (US), current annual losses attributed to this infectious disease amount to approximately 664 million US dollars [2]. The global dissemination of PRRSV is a prime example of how modern agricultural trade networks can facilitate the rapid, long-distance spread of high-consequence livestock pathogens [3,4]. While PRRSV itself is not zoonotic, its profound economic impact threatens food security, which has direct consequences for human health and welfare. The effect of the global live swine trade on the spread of viral infections was evident during the 2009 H1N1 pandemic, which highlighted the critical role of international trade in the global ecology and evolution of swine viruses [5].

Furthermore, elucidating viral dissemination pathways like those explored here provides a critical model for understanding the introduction and spread of zoonotic pathogens that exploit the same global trade routes. The reproductive form of PRRSV-2 is characterized by symptoms such as anorexia, late-term abortions occurring between days 107-112 of gestation, an increase in the number of stillborns, mummified, and weak-born piglets, higher pre-weaning mortality, and a delayed return to estrus. Pronounced hyperpnea, fever, and interstitial pneumonitis occur in young pigs with this disease, and mild flu-like symptoms are seen in nursing, growing, and finishing pigs [6].

In August 1995, an outbreak of severe respiratory illness and increased mortality among weaned pigs on the CA-CART farm in Costa Rica’s Cartago province led to the country’s first diagnosis of PRRSV. Necropsies performed on the affected piglets showed a range of symptoms, including respiratory lesions, pleuritis, peritonitis, arthritis, and skin lesions that developed into scabs. A retrospective analysis of farm mortality data (1990-1995) did not find a seasonal pattern but did reveal annual variations. Crucially, this analysis demonstrated that piglets from gilts imported from the United States experienced a 10.65% average mortality rate, significantly higher than the 8.11% rate observed in piglets from non-imported mothers (p = 0.002). Initially, the so-called “mystery swine disease,” as it was known at the time, was linked to the Swine Infertility and Respiratory Syndrome (SIRS) virus, specifically the ATCC VR-2332 strain in North America and the Lelystad virus strain in Europe.[7]. Subsequently, the International Committee on Taxonomy of Viruses (ICTV) classified these viruses under the Arteriviridae family, genus Porartevirus, identifying them as two different species: PRRSV-1 (previously known as European, EU Lelystad virus) and PRRSV-2 (previously known as North American, NA VR-2332 strain [8].

The *Porartevirus* genus is characterized by enveloped viruses with a single-stranded, positive-sense RNA genome. The genome is approximately 15 kb long and consists of 10 open reading frames (ORFs). Three-quarters of the PRRSV genome consists of ORF1a and ORF1b, which encode 14 nonstructural proteins. The terminal part consists of eight partially overlapping ORFs that encode the structural proteins that constitute the envelope (ORF2a, ORF3, ORF4, ORF5 5 and ORF5a), matrix (ORF6), and nucleocapsid (ORF7).

Glycoprotein 5 (GP5), encoded by ORF5, is the most widely studied PRRSV protein due to its high genetic diversity and its role in virulence, virus-host interaction, and immunity [9–11]. PRRSV-2 has been classified into nine distinct lineages based on a comprehensive collection of ORF5 sequences [4] and grouped in two clades denoted as lineages 1–2 and lineages 6–9, respectively [12]

From August 25, 2015, to May 11, 2016, we conducted surveillance for swine disease in 25 farms distributed across all 7 provinces of Costa Rica (Alajuela, Cartago, Guanacaste, Heredia, Limón, Puntarenas, and San José). The farms were categorized as large (8), medium (12), and small (5), and depending on the farm’s size, 40 to 70 serum samples were collected [13]. The results of this study demonstrated that during the surveillance period, 44% (11 out of 25) of the farms tested positive for PRRSV-2, with these farms spread across six of the seven provinces of Costa Rica. Additionally, 58% (344 out of 596) of the pigs sampled were found to be seropositive [14]. Sequencing and phylogenetic modeling of ORF5 of the collected samples grouped the sequences analyzed into three clusters distributed in two lineages. Viral sequences from San Jose, Alajuela, Puntarenas, and Heredia belonged to Lineage 5, while samples from the Cartago farm clustered in a separate clade related to Lineage 1 [15].

Here, we describe the recent evolutionary history of PRRSV-2 in Costa Rica using Bayesian phylodynamic and phylogeographic modeling of partial ORF5 gene sequences previously described [15] and generated during this project. We aim to shed light on the transmission dynamics of the virus in the country to better inform control strategies.

## MATERIALS AND METHODS

### Farms Studied

#### Case History and Sampling Periods

During the period between July 2015 and July 2016, as part of a national serological study[15], 72 serum samples were collected from piglets aged 1 to 15 weeks on this farm.

From March to April 2019, ninety samples were collected to be amplified and sequenced from nine farms, including the CA−CART farm, specifically for the purpose of identifying and sequencing PRRSV [15]. Nine samples were also collected in August 2021 on the CA−CART farm as part of this project.

Figure S1 shows the locations where the samples analyzed in this study were collected.

In June 2021, the Nebraska strain, a modified live PRRSV-2 vaccine, was introduced on the CA−CART farm. The same reproductive and production parameters were monitored to evaluate the vaccine’s effects. In August 2024, after 38 months of vaccination, five blood samples from nursery pigs were collected to detect the presence of PRRSV using qRT-PCR and cell culture.

#### Statistical Analysis

Statistical analyses were performed with the software package EPI INFO version 6.03 (CDC 1600 Clifton Road, Atlanta, GA 30329-4027 USA). To calculate the percent average mortality, the number of animals that died each month was divided by the weaning population at the beginning of the month minus the weaning population at the end of the month and multiplied by 100. We also investigated potential risk factors for mortality, including weather, season, month, and year of death, and pig importation history. The odds ratio was calculated using Fisher’s exact test and the Yates-corrected chi-square on single 2-by-2 contingency tables.

#### Phylogenetic analyses using maximum likelihood and Bayesian approaches

We analyzed a total of 10 ORF5 sequences generated in Costa Rica: 8 sequences isolated between March and April 2019 [15] and 2 sequences collected between February and August 2021, which were then sequenced using the Sanger method as previously described [15].

To compile a genetic background dataset, we downloaded ORF5 PRRSV sequences from GenBank in August 2023 with more than 89% identity compared to the 10 Costa Rican sequences using BLAST [16].. A total of 10 sequences from Costa Rica and 92 sequences from other countries were aligned using Bioedit v 7.2.5 [17]. The sequences were analyzed by IQ-Tree v1.6.2 [18] to determine the best substitution model and discard identical sequences. The stability of the obtained tree topology was assessed by bootstrapping (1000 replicates). TempEst v 1.5.3 [19] was used to evaluate the temporal signal, enabling the identification and exclusion of any outlier sequences and also to estimate the rate of evolution in substitutions per site per year. RDP4 was employed to determine recombination events in the ORF5 genetic dataset [20]. The final dataset comprised 10 sequences from Costa Rica and 70 sequences generated worldwide (Supplementary Table S1). We employed a Bayesian phylogenetic framework in the BEAST v1.10.4 software package [21] to reconstruct time-calibrated phylogenetic and phylogeographic histories. A GTR+F+I+G4 substitution model, an uncorrelated relaxed clock [22], and the Bayesian SkyGrid coalescent prior were applied. To accommodate discrete variables in ORF5 datasets, we used an asymmetric substitution model with Bayesian Stochastic Search Variable Selection (BSSVS). Six independent Markov chain Monte Carlo (MCMC) chains were run for 50 million generations each [23]. The 6 replicates were combined using LogCombiner v1.10.4, after confirming convergence using Tracer v1.7.2.

The trees were logged every 50,000^th^ MCMC step, and the tree distribution was summarized using TreeAnnotator v1.10.4 as a maximum clade credibility (MCC) tree, with median heights as the node heights in the tree. FigTree v1.4.3 was used to visualize the phylogenetic trees (http://tree.bio.ed.ac.uk/software/figtree/). The Bayes Factor summaries were obtained using spreadviz.org, and the transmission dynamics were visualized using ggmap [24].

We determined the PRRSV lineages for each sequence in the MCC tree using a dual approach. First, we applied the Nextclade nomenclature system [25,26], and then we used the Sequence Demarcation Tool (SDT) to classify the sequences based on pairwise genetic identity [26–28]

## RESULTS

During the period between July 2015 and July 2016, a prospective study was conducted to determine the overall prevalence of PRRSV in Costa Rica. Two random samplings established a 26.9% of the animals sampling (344/1281), 95% CI (24.5–29.4), were positive to PRRSV, while the median within-herd seroprevalence in the seropositive farms was 58%, 344 seropositive in 596 pigs sampled [14]. In the CA−CART farm, 62 out of 72 serum samples (86.1%) tested positive.

Between March and April 2019, a follow-up sampling was conducted as an extension of the previous study to confirm whether PRRSV-2 was the sole strain circulating in Costa Rica. Of the 90 samples collected in 9 PRRSV-positive farms, 27 yielded partial sequences. This study confirmed the presence of PRRSV-2 in Costa Rica, with no evidence of PRRSV-1 detected [14], leading to the approval of PRRSV-2 vaccines in the country.

In July 2020, a severe outbreak occurred on the CA−CART farm, resulting in a significant rise in mortality, 24% in the farrowing stage, 15% in the nursery stage, and increasing to 24% in the nursery and 12% in the growing phases by August.

Consequently, in June 2021, a modified live attenuated Nebraska strain PRRSV-2 vaccine was administered on the CA−CART farm. The effects of this vaccination are summarized in Table 1.

**Table 1:**
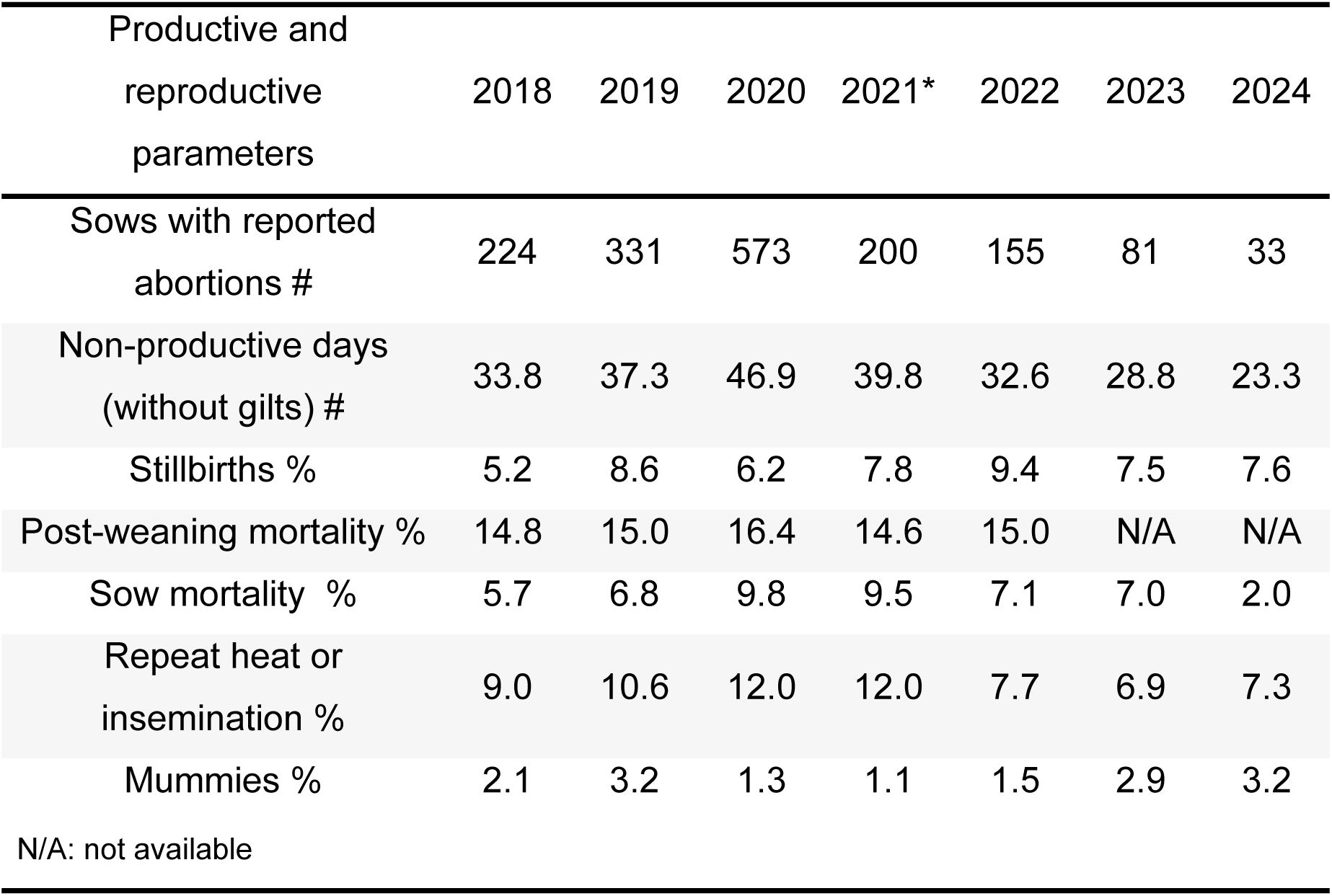
Comparison of productive and reproductive parameters before and after the use of a PRRSV-2 vaccine between 2018 and 2024. *Introduction of the vaccine in June 2021. Note the sharp decline in sow abortions and mortality and the reduction in non-productive days in the post-vaccination years (2022-2024) compared to the pre-vaccination peak during the 2020 outbreak.

| Productive and reproductive parameters | 2018 | 2019 | 2020 | 2021* | 2022 | 2023 | 2024 |
| --- | --- | --- | --- | --- | --- | --- | --- |
| Sows with reported abortions # | 224 | 331 | 573 | 200 | 155 | 81 | 33 |
| Non-productive days (without gilts) # | 33.8 | 37.3 | 46.9 | 39.8 | 32.6 | 28.8 | 23.3 |
| Stillbirths % | 5.2 | 8.6 | 6.2 | 7.8 | 9.4 | 7.5 | 7.6 |
| Post-weaning mortality % | 14.8 | 15.0 | 16.4 | 14.6 | 15.0 | N/A | N/A |
| Sow mortality % | 5.7 | 6.8 | 9.8 | 9.5 | 7.1 | 7.0 | 2.0 |
| Repeat heat or insemination % | 9.0 | 10.6 | 12.0 | 12.0 | 7.7 | 6.9 | 7.3 |
| Mummies % | 2.1 | 3.2 | 1.3 | 1.1 | 1.5 | 2.9 | 3.2 |

N/A: not available

In August 2021, all nine sera collected from the farm CA−CART tested positive for PRRSV-2 by qRT-PCR, and one of these samples was successfully sequenced. In 2024, five additional serum samples were taken from nursery piglets on the same farm. Two of them tested positive for PRRSV-2 by qRT–PCR; however, due to low viral loads, sequencing and viral isolation were not possible.

### Genomic epidemiology analysis

After the sequences were analyzed by IQ-Tree v1.6.12, 22 duplicate sequences were removed. Among the remaining 80 sequences, 6 sequences exhibited recombination events as identified by the RDP4 program: three originated from the US, two from India, and one from China. Details on these sequences, including their major and minor parents and the recombination breakpoints, are shown in Table S2. The final phylogenetic tree shows the distribution of the Costa Rican sequences, which group into three distinct clusters: 1CRC, 3CRC, and 4CRC. Cluster 1CRC (red) includes sequences collected in three farms located in the provinces of San Jose, Puntarenas, and Alajuela, cluster 3CRC (orange) contains sequences from two farms in Cartago (including CA-CART) and Heredia, while cluster 4CRC (yellow) is comprised of sequences collected from a farm in Heredia (Figure 1).

**Figure 1.**
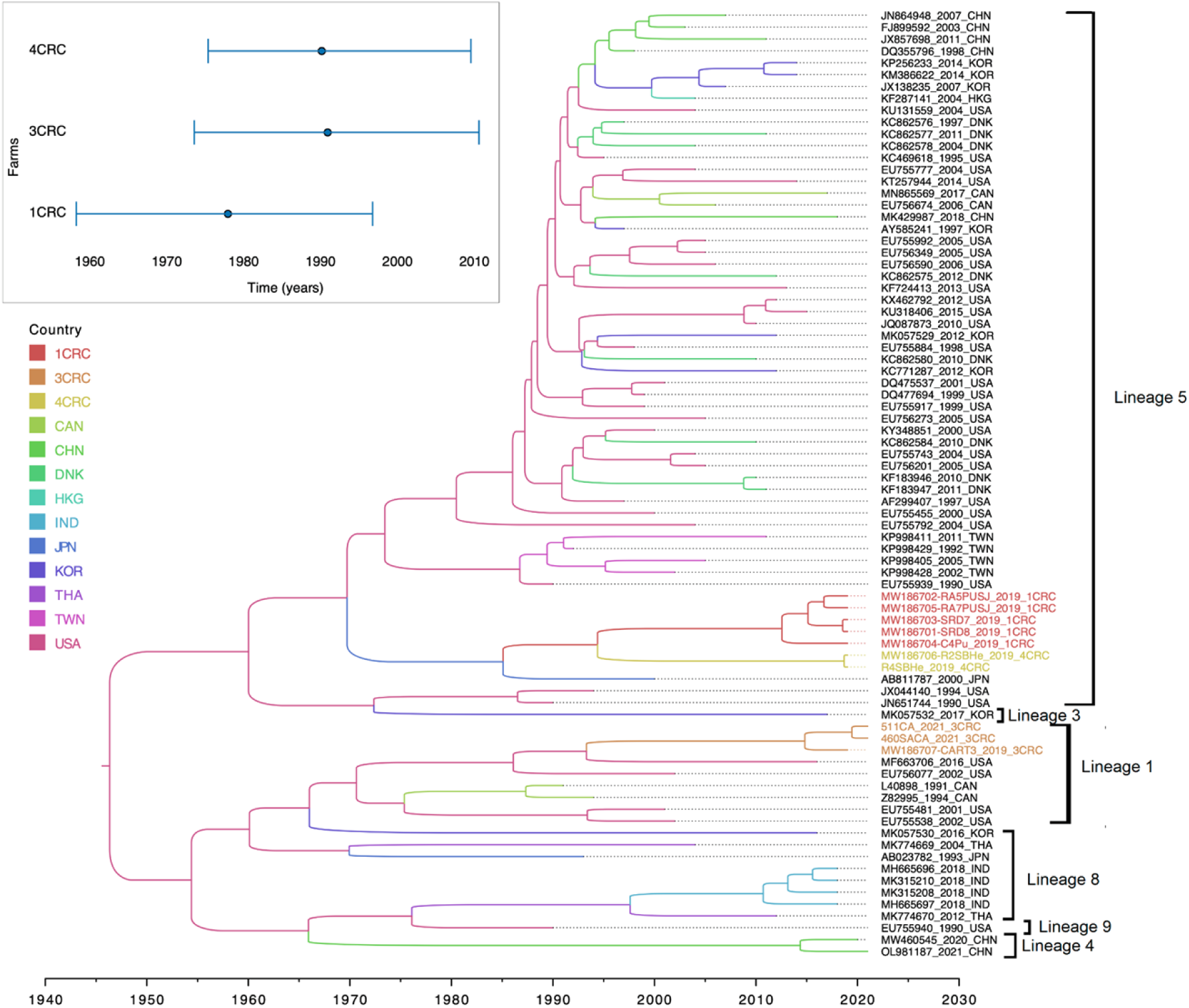
Phylogeographic tree of the reconstructed PRRSV transmission dynamics. Maximum clade credibility tree with branches colored by country estimated as the most probable trait. Inset: Estimated time to the most recent common ancestor (TMRCA and 95% highest posterior density interval) of PRRSV-2 farm clusters.

The Time to Most Recent Common Ancestor (TMRCA) of the 1CRC sequences was estimated at 1978.72 (95% highest posterior density (HPD) interval = 1959.01-1997.54) with a 0.63 probability of having originated in Japan (posterior probability 0.98), though this may reflect earlier swine importations from the US to Japan. For cluster 3CRC, the TMRCA was 1991.69 (95% HPD interval = 1974.34-2011.34) with a 0.96 probability of originating in the US (posterior probability: 0.85).

Evidence suggests that the introduction of the 1CRC lineage into Costa Rica may have led to the emergence of cluster 4CRC (location probability 0.30; posterior probability 0.99). The TMRCA of the 4CRC was estimated to be 1990.24 (95% HPD interval = 1976.15-2010.30). The phylogenetic tree shows that the Costa Rican sequences are distributed across two lineages: lineage 1, which includes the three Costa Rican 3CRC sequences, and lineage 5, which contains the 1CRC and 4CRC sequences [15] (Figure 1). The average pairwise genetic distance (% difference) for six of the nine intra- and inter-sublineages described by Shi et al. (2010) is presented in the supplementary Figure S2, as well as the average pairwise genetic distances of the Costa Rican sequences. Including the Costa Rican sequences, the intra-lineage similarity within L1 is 91% and 96% within L5. The genetic distance between L1-3CRC and other L1 sequences is 11%, while the distances between the L5-1CRC and L5-4CRC groups are 11% and 10%, respectively. The genetic distance between lineages L1 and L5 is also 11%, highlighting a comparable level of divergence to that observed between the Costa Rican subgroups and their assigned lineages. The lineage classification according to Nextclade is presented in Supplementary Table S3.

### Population dynamics of PRRSV

The demographic plot of PRRSV-2 (Figure 2) shows the growth of the viral effective population size (Ne) between 1992 and 2000 and from 2000 – 2008 in Costa Rica. The curve shows a decrease in the viral diversity from 2008 to 2020.

**Figure 2:**
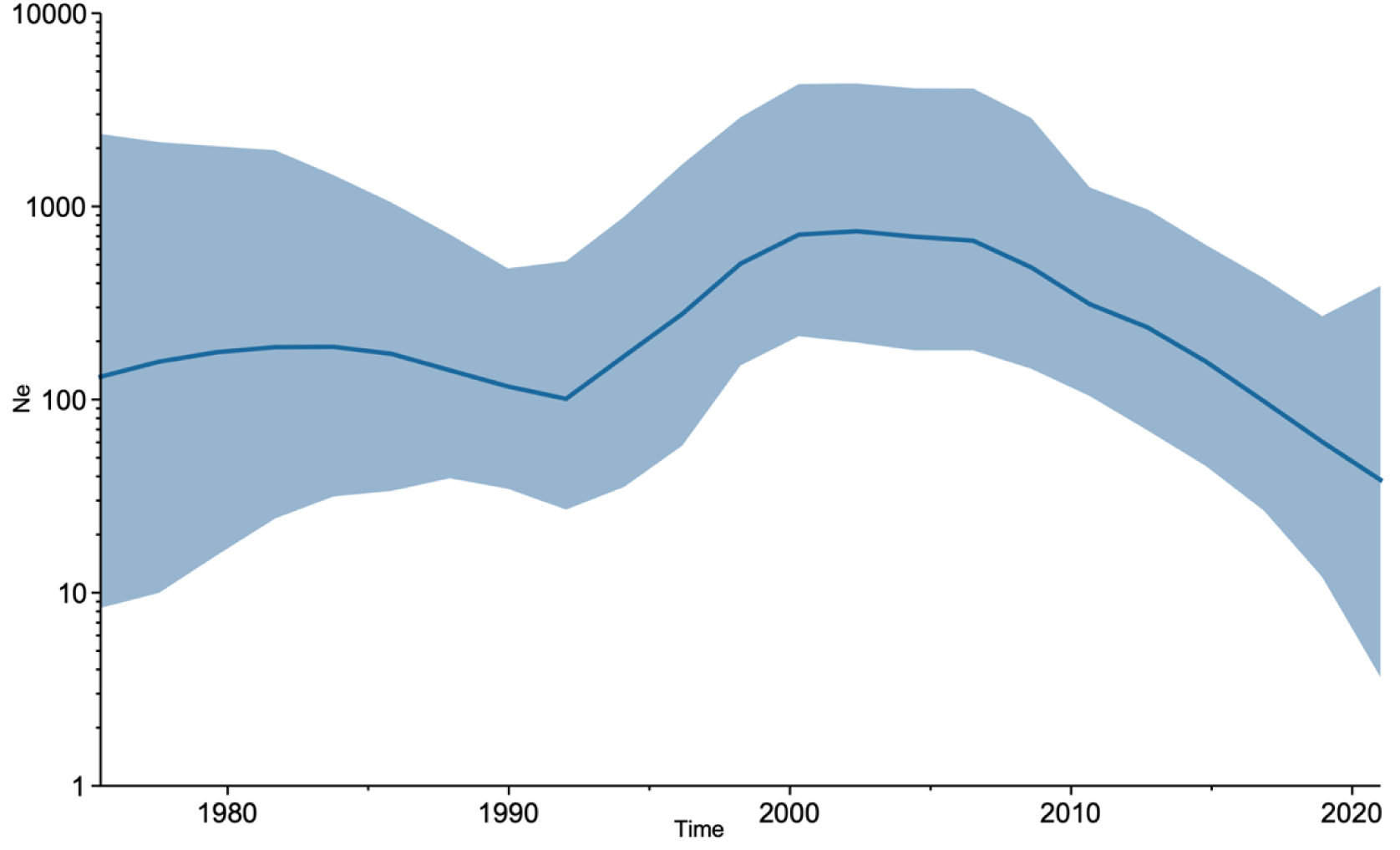
Viral population dynamics of PRRSV. Bayesian Skygrid plot where the y-axis indicates the effective population (Ne) and the x-axis shows the time expressed in years. The solid line in the graph indicates the median Ne, while the blue area indicates the 95% highest posterior density interval.

### Phylogeographic reconstruction

Our analysis identified three independent introductions of PRRSV-2 into Costa Rica. Approximately 50% of the genomic sequences from Costa Rica (cluster 1CRC) appear to have originated from a single introductory event from Japan (Figure 3), supported by a Bayes factor (BF) value for the node of 9.92. We estimated that 30% of the viral sequences originated from the United States (cluster 3CRC) with a BF value of 5.88. Within Costa Rica, evidence suggests that the virus diverged from cluster 1CRC to form cluster 4CRC between January 1976 and March 2010, supported by a BF value of 6.62 (Figure 3).

**Figure 3.**
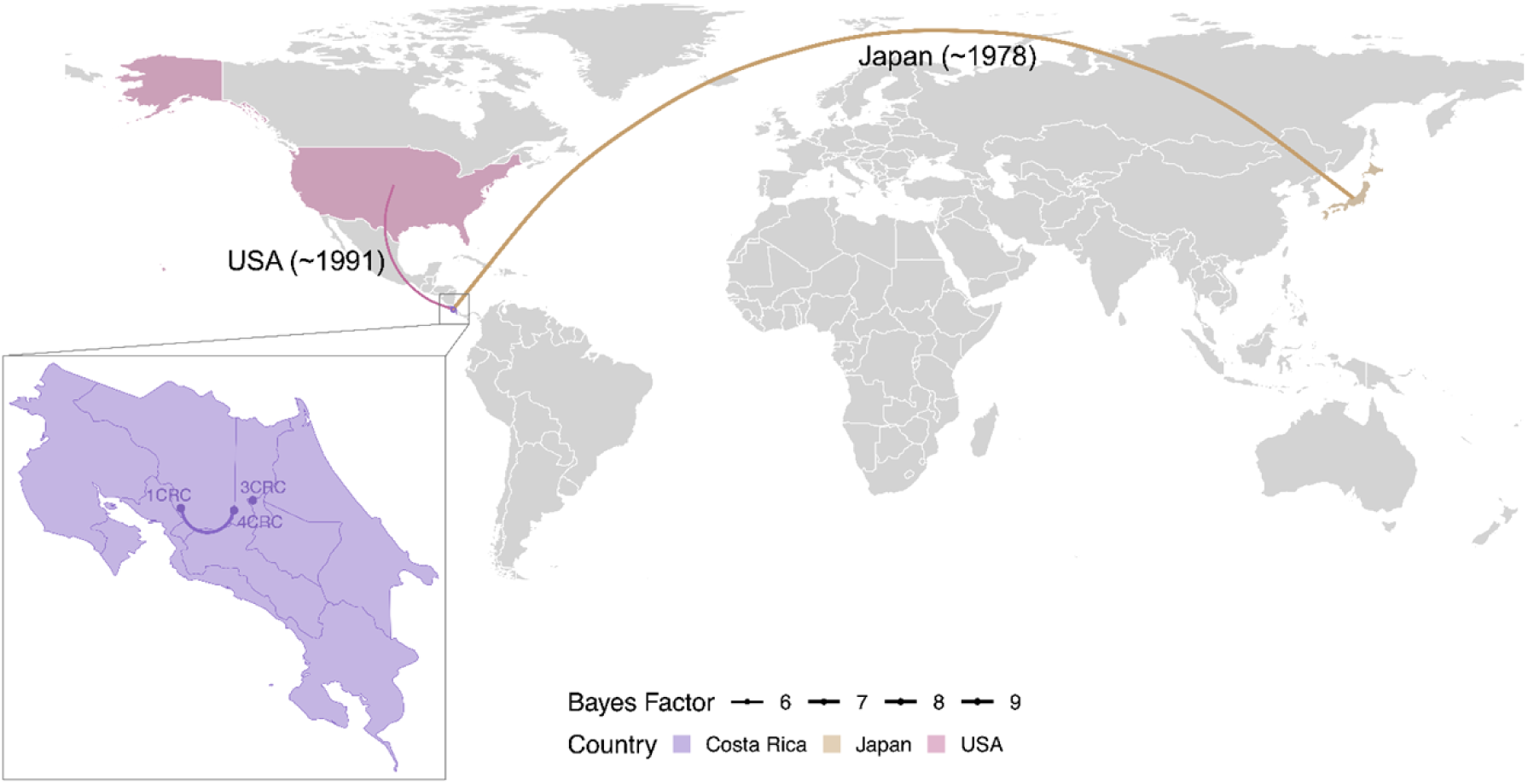
Tracing the origin of PRRSV-2 introduction into Costa Rica. The concave arcs indicate the westward transmission events, and convex arcs indicate eastward transmission events between locations, which represent branches in the MCC tree. The thickness of the arc is based on Bayes factors, and the color depends on the originating country.

## DISCUSSION

Since its emergence in the late 1980s, PRRSV-2 has spread rapidly and become enzootic in pig populations worldwide. Although PRRSV-2 is not a zoonotic disease, it has a significant impact on the swine industry. The economic cost of an acute PRRSV-2 outbreak in breeding herds was estimated at USD 255 per sow and USD 6.25 to USD 15.25 per growing pig [29]. In the US alone, the annual cost of PRRSV outbreaks to the US swine industry is estimated at around USD 560 million [30]. Economic losses during the breeding and farrowing phases are primarily due to a reduction in the number of weaned pigs and reduced farrowing rates, accounting for 11.9% of the total annual economic loss. In the nursery and grower-finisher phase, losses are attributed to the increased mortality, decreased feed efficiency, and lower average daily weight gain, accounting for 35.9% and 52.2%, respectively [31]

Following the initial diagnosis of PRRS in Costa Rica, 7 out of 8 pigs randomly sampled from the CA−CART farm in March 1996 tested positive for PRRSV-2 antibodies, corresponding to a seroprevalence of 87.5%. Two decades later, this seroprevalence remained significantly higher than the national average reported by Meléndez et al. (2021), which was 26.9% (344/1,281) among pigs sampled across 25 farms (95% CI: 24.5–29.4). Notably, during the Meléndez study, the seroprevalence at the CA−CART farm was 86.1% (62/72), also exceeding the median within-herd seroprevalence observed in seropositive farms, which was 58% (344/596). These findings suggest that PRRSV seroprevalence at the CA−CART farm has remained consistently high over two decades, indicating that the control measures implemented have been ineffective and that persistent viral circulation continues within the herd.

Our results showed that peaks of PRRSV-2 infection globally were also observed in Costa Rica in 2000 and 2020. A significant decrease was observed only after the introduction of the vaccine in 2021, leading to improvements in reproductive and production indices on the CA-CART farm. Moreover, a reduction in viral transmission of the virus was evident, as illustrated in Table 1. The benefits of vaccination have also been reported in other studies comparing different vaccination strategies, such as vaccinating only sows or vaccinating both sows and piglets [32,33]

Previous studies have shown that the nucleotide variability of the PRRSV ORF5 gene enhances the resolution of phylogenetic relationships[12,15,34]. This variability makes possible the classification of PRRSV-2 sequences into 9 lineages, which have ∼10% nucleotide divergence [28]

The genetic diversity across PRRSV ORFs indicates that the virus evolves via mechanisms beyond the simple accumulation of random mutations. Statistical analyses of the nucleotide sequences have provided strong evidence for intragenic recombination or gene conversion in ORFs 2, 3, 4, 5, and 7, but not in ORF 6 [35]. In our analysis, we detected six recombinant sequences. Recombination in ORF 5 has been reported between lineages 1 and 2 with common recombination breakpoints at nucleotide positions 126, 350, and approximately 430 [4]. In our dataset, similar breakpoint locations were observed in background dataset sequences; however, recombination events were not detected among the Costa Rican sequences.

Since the emergence of PRRSV-2 in the US in the late 1980’s in the US, 20-40% of breeding herds have experienced outbreaks annually. In Costa Rica, the first laboratory confirmed case was reported on March 15, 1996, although the initial clinical cases of this outbreak occurred in August 1995. The first importation of pigs from the US to the affected farm was recorded in 1988, with the third in 1992. Notably, increased mortalities were recorded during the nursery stage in both 1991 and 1993.

Bayesian phylogeographic analysis was used to explore the geographic dispersion of PRRSV-2 within Costa Rica. ORF5 gene sequences, isolated between 2019-2021 in Costa Rica, were compared with publicly available international sequences. Phylodynamic modeling estimated the TMRCA for the 3CRC cluster, comprising PRRSV-2 sequences collected from the CA-CART farm during 2019-2021 and a sequence collected in the US in 2002, tracing back to an ancestor in the US in 1991.69. For the 1CRC and 4CRC clusters, the TMRCAs were estimated at 1978.72 and 1990.92, respectively. Our phylodynamic estimate for the introduction of the US-originating 3CRC cluster (1991.69) aligns remarkably well with farm records of increased nursery-stage mortality in 1991 and 1993, which followed pig importations from the US. This provides strong, independent evidence that these importations were the likely source of this lineage, and align with historical records of pig importation from the US to these farms in 1986. However, there is no information on whether animals were imported from outside Costa Rica before 1986.

Our phylodynamic modeling not only identified the likely origins of PRRSV-2 in Costa Rica but also provided a precise timeline that aligned remarkably with historical farm records of increased mortality following swine importations. The successful reconstruction of these decades-old introductions underscores the power of phylodynamics to elucidate complex transmission pathways long after a pathogen’s initial emergence.

In a previous study, it was determined that the viral sequences from 1CRC and 4CR belong to lineage 5 while those from group 3CRC are part of lineage 1 [15]. It was suggested that 99.2% of the Canadian sequences from lineage 5 belonged to the modified live vaccine (MLV)-related sublineage 5.1 [4]. According to Shi et al, in the early 2000’s, a significant shift in the genetic composition of non-vaccine-related PRRSV-2 occurred in the Midwest regions of the US, where lineages 6–9 sequences used to dominate and lineages 1–2 were rare. After 2001, the sampling frequency of lineages 1–2 gradually increased. The trend continued uninterrupted, and as a result, 77% of the most recent samples collected from the US Midwest region belong to lineages 1–2 [4]. The percentage of genetic divergence observed between the Costa Rican sequences and their respective lineages may be influenced by the limited number of sequences analyzed in this study. According to Nextclade, the 1CRC and 4CRC sequences are classified not only within lineage 5 but more specifically within the sublineage 5A, while two sequences from the 3CRC group are classified as lineage 1, sublineage E.

Interestingly, sequence MW186707-CART3_2019_3CRC is classified as L4 by Nextclade, which is inconsistent with its placement within lineage L1 based on the reconstructed MCC tree topology shown in Figure 1. This is likely due to its short length of only 253 nucleotides. Nextclade determines lineage assignments based on the mutations separating the query sequence from its closest reference match (Aksamentov et al., 2021). However, when the sequences were translated into amino acids (Figure S3), no shared mutations were identified between MW186707-CART3_2019_3CRC and the L4 reference sequences (MW460545 and OL981187), suggesting that the classification may be unreliable in cases of limited sequence length and potential quality issues.

Phylogenetically, viruses 1CRC and 4CRC (lineage 5) cluster closely with strains from North America and East Asia. This relationship reflects the widespread distribution of lineage 5 due to its association with vaccine strains. However, characterizing the transmission network of vaccine-associated lineages is challenging because (i) it is extremely difficult to distinguish vaccine-derived strains from field viruses and (ii) the dynamics of vaccine-derived viruses differ substantially from those of field viruses [4]. Lineage 1 viruses were initially detected in Canada and only later in the US [4]; however, our phylodynamic modeling suggests that the virus may have been circulating in the US for decades before its detection.

The SkyGrid plot (Figure 2) provides valuable insights into the temporal dynamics of PRRSV-2 transmission. The initial period between 1992 and 2000 witnessed a significant increase in the effective population size (Ne), suggesting a rapid expansion of the virus. From 2000 to 2008, the effective population size (Ne) declined, indicating a possible decrease in viral transmission or a stabilization of the epidemic. The outbreak reported in 2000 on the CA-CART farm led to a drastic reduction in the number of sows sent to slaughter. On a national level, this decrease could be attributed to factors such as improved biosecurity measures adopted in farms [36,37] or changes in swine husbandry practices. The subsequent period, from 2008 to 2020, showed further reduction in Ne, suggesting a continued decline in viral transmission or a shift toward a more stable endemic state. This trend could be due to the introduction of the vaccine [38].

The investigation of PRRSV-2 introduction and diversification in Costa Rica provides valuable insights into the global spread and evolution of this virus. Our analysis identified two independent introductions from Japan and the United States, with a significant proportion of Costa Rican sequences clustering with the Japanese virus. This suggests that the initial introduction from Japan played a major role in shaping the genetic diversity of PRRSV-2 in Costa Rica. However, it is possible that the viral sequence collected in Japan was sampled from pigs bought from the US in the mid-1980s [28]. The emergence of the 4CRC cluster in Costa Rica indicates the potential for local transmission and adaptation of the virus. However, the limited number of sequences available for analysis may have influenced the resolution of the phylogenetic tree and the precise timing of diversification events. It is important to note that the BF values associated with the inferred introductions and diversification events provide moderate to strong statistical support. Further studies with a larger sample size could provide more detailed insights into the evolutionary history of PRRSV-2 in Costa Rica.

Our phylodynamic modeling indicates that pigs imported from the US likely seeded the initial cases of PRRSV-2 in Costa Rica. These animals were subsequently distributed to local farms, facilitating the spread of the virus within the Costa Rican swine population. For farms clustered in the 1CRC group, which included sequences from three different provinces: San Jose, Puntarenas and Alajuela, virus dispersal was likely influenced by hog transportation within multi-site production systems [4]. It is clear that PRRSV-2 is still an important threat for the swine industry in Costa Rica, but it could be controlled through the implementation of strict biosecurity measures. Several methods for controlling or eradicating the disease have been described worldwide [31]. However, despite the implementation of these control methods and strict adherence to biosecurity procedures, many herds continue to experience reinfection due to the persistence of the virus in the area.

As a result, some producers and veterinarians are considering the implementation of a voluntary regional program that would include all herds within a specific area. For such a program to be effective, control strategies, such as gilt acclimation, herd closure, depopulation-repopulation, and others, must be applied consistently across all infected farms within the region [31,39]. It is critical to have the local producers’ acceptance and cooperation; communication and education, including this study’s findings, are important tools to increase producers’ awareness and compliance.

Beyond the local importance for Costa Rican swine health, our findings provide a textbook case study on the value of integrated phylodynamic modeling for safeguarding both animal and human health. Although PRRSV is not zoonotic, its economic impact underscores how animal diseases can affect human welfare through threats to food security and economic stability. More importantly, we demonstrate how reconstruction of the viral evolutionary history and spatio-temporal dynamics can unveil complex introduction histories, a framework directly applicable to tracing the emergence of zoonotic viruses like Nipah [40] or swine influenza [5] that travel through the same global livestock trade networks. This integrated approach can therefore be applied to other high-consequence pathogens to inform trade policies and enhance global health security.

This report aims to shed light on the routes of PRRSV-2 introduction into Costa Rica and describe the epidemiological features leading up to the first confirmed diagnosis of PRRS in the country. The limited number of available sequences prevented the identification of a closer TMRCA between Costa Rican and US sequences. This likely explains why a sequence from Japan appears as the most probable source of the virus introduced into the farms associated with the 1CRC and 4CRC clusters. Another limitation of the study was the inability to obtain additional sequences from these farms. Following the introduction of the vaccine, low viremia levels hindered sequencing efforts, making it impossible to determine whether the vaccine strain had replaced the original lineage 1 virus.

Almost 30 years after the first diagnosed case of PRRSV-2 in Costa Rica, PRRS continues to be a serious problem for pig producers. This underscores the importance of understanding the viral dynamics that drive epidemics to protect animal agriculture and, by extension, human economic well-being and food security. The methods and findings presented here not only inform control measures for PRRSV but also enhance our preparedness for future zoonotic threats that may emerge through identical pathways.

## DATA AVAILABILITY

Viral sequences are available in GenBank with accession numbers PV665664-PV665665.

## AUTHOR CONTRIBUTIONS

Conceptualization: BL, NST; Methodology: NST; Formal analysis: BL, ST, SK, NST; Investigation: BL; Resources: BL; Data Curation: BL, ST, SK, NST; Writing - Original Draft: BL and ST; Writing - Review & Editing: BL, NST, DS; Visualization: SK, NST; Supervision: NST; Project administration: NST; Funding acquisition: BL.

## ACKNOWLEDGEMENTS

We are grateful for the essential contributions of our collaborators. We thank the SENASA staff for collecting the field samples. The opinions expressed in this article are those of the authors and do not reflect the views of the National Institutes of Health, the Department of Health and Human Services, the United States government, or the National Animal Health Service, SENASA, Costa Rica.

## Supplementary Material

### Supplementary Tables

**Table S1.** Sequences analyzed in this study.

| GenBank ID | Collection year | Country |
| --- | --- | --- |
| AB023782 | 1993 | JPN |
| AB811787 | 2000 | JPN |
| AF299407 | 1997 | USA |
| AY585241 | 1997 | KOR |
| DQ355796 | 1998 | CHN |
| DQ475537 | 2001 | USA |
| DQ477694 | 1999 | USA |
| EU755455 | 2000 | USA |
| EU755481 | 2001 | USA |
| EU755538 | 2002 | USA |
| EU755743 | 2004 | USA |
| EU755777 | 2004 | USA |
| EU755792 | 2004 | USA |
| EU755884 | 1998 | USA |
| EU755917 | 1999 | USA |
| EU755939 | 1990 | USA |
| EU755940 | 1990 | USA |
| EU755992 | 2005 | USA |
| EU756077 | 2002 | USA |
| EU756201 | 2005 | USA |
| EU756273 | 2005 | USA |
| EU756349 | 2005 | USA |
| EU756590 | 2006 | USA |
| EU756674 | 2006 | CAN |
| FJ899592 | 2003 | CHN |
| JN651744 | 1990 | USA |
| JN864948 | 2007 | CHN |
| JQ087873 | 2010 | USA |
| JX044140 | 1994 | USA |
| JX138235 | 2007 | KOR |
| JX857698 | 2011 | CHN |
| KC469618 | 1995 | USA |
| KC771287 | 2012 | KOR |
| KC862575 | 2012 | DNK |
| KC862576 | 1997 | DNK |
| KC862577 | 2011 | DNK |
| KC862578 | 2004 | DNK |
| KC862580 | 2010 | DNK |
| KC862584 | 2010 | DNK |
| KF183946 | 2010 | DNK |
| KF183947 | 2011 | DNK |
| KF287141 | 2004 | HKG |
| KF724413 | 2013 | USA |
| KM386622 | 2014 | KOR |
| KP256233 | 2014 | KOR |
| KP998405 | 2005 | TWN |
| KP998411 | 2011 | TWN |
| KP998428 | 2002 | TWN |
| KP998429 | 1992 | TWN |
| KT257944 | 2014 | USA |
| KU131559 | 2004 | USA |
| KU318406 | 2015 | USA |
| KX462792 | 2012 | USA |
| KY348851 | 2000 | USA |
| L40898 | 1991 | CAN |
| MF663706 | 2016 | USA |
| MH665696 | 2018 | IND |
| MH665697 | 2018 | IND |
| MK057529 | 2012 | KOR |
| MK057530 | 2016 | KOR |
| MK057532 | 2017 | KOR |
| MK315208 | 2018 | IND |
| MK315210 | 2018 | IND |
| MK429987 | 2018 | CHN |
| MK774669 | 2004 | THA |
| MK774670 | 2012 | THA |
| MN865569 | 2017 | CAN |
| MW460545 | 2020 | CHN |
| OL981187 | 2021 | CHN |
| Z82995 | 1994 | CAN |
| MW186707-CART3 | 2019 | 3CRC |
| MW186701-SRD8 | 2019 | 1CRC |
| MW186702-RA5PUSJ | 2019 | 1CRC |
| MW186703-SRD7 | 2019 | 1CRC |
| MW186704-C4Pu | 2019 | 1CRC |
| MW186705-RA7PUSJ | 2019 | 1CRC |
| MW186706-R2SBHe | 2019 | 4CRC |
| R4SBHe | 2019 | 4CRC |
| 511CA | 2021 | 3CRC |
| 460SACA | 2021 | 3CRC |

**Table S2.**
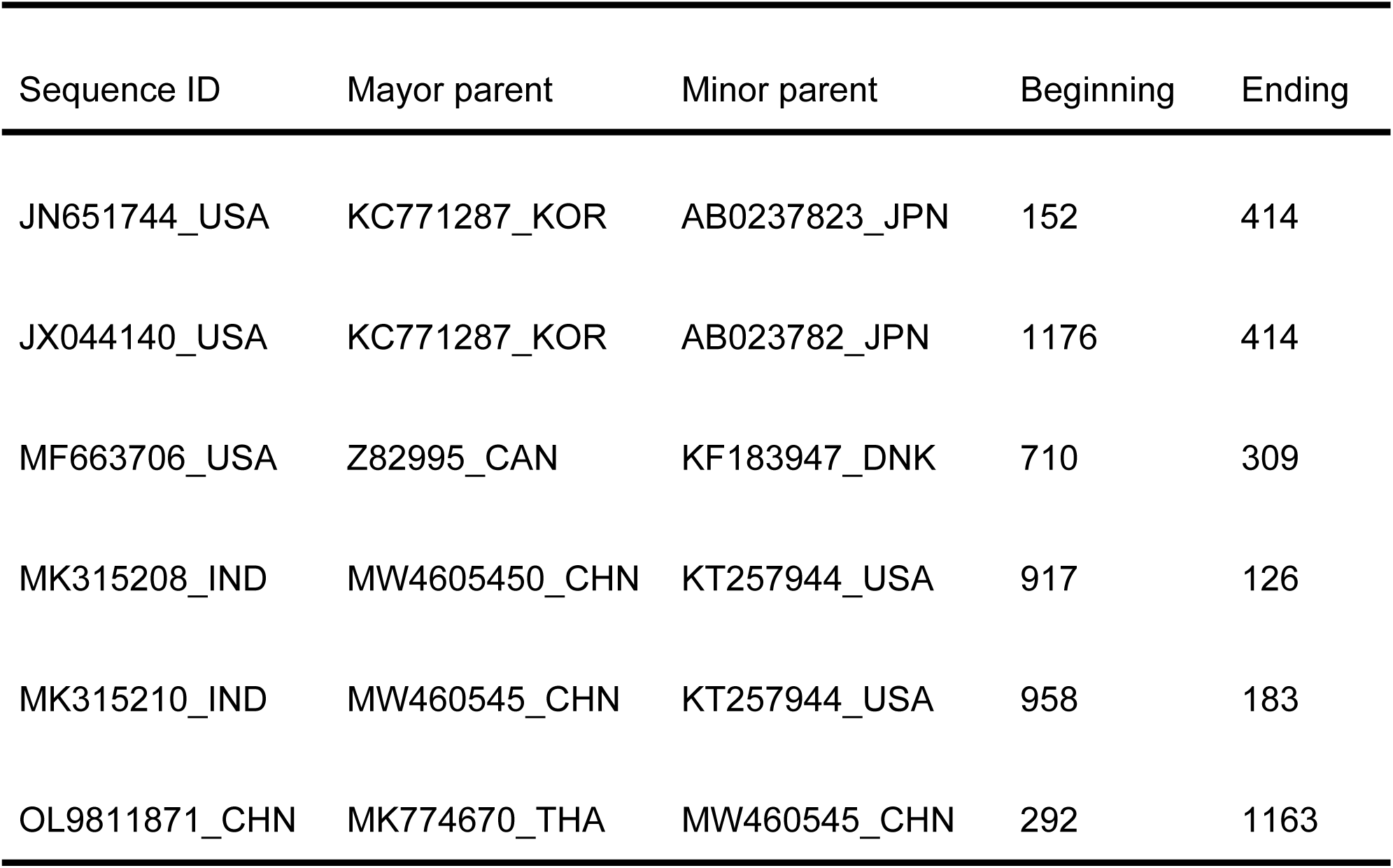
Recombination events detected in the PRRSV-2 ORF5 sequences.

| Sequence ID | Mayor parent | Minor parent | Beginning | Ending |
| --- | --- | --- | --- | --- |
| JN651744_USA | KC771287_KOR | AB0237823_JPN | 152 | 414 |
| JX044140_USA | KC771287_KOR | AB023782_JPN | 1176 | 414 |
| MF663706_USA | Z82995_CAN | KF183947_DNK | 710 | 309 |
| MK315208_IND | MW4605450_CHN | KT257944_USA | 917 | 126 |
| MK315210_IND | MW460545_CHN | KT257944_USA | 958 | 183 |
| OL9811871_CHN | MK774670_THA | MW460545_CHN | 292 | 1163 |

**Table S3.**
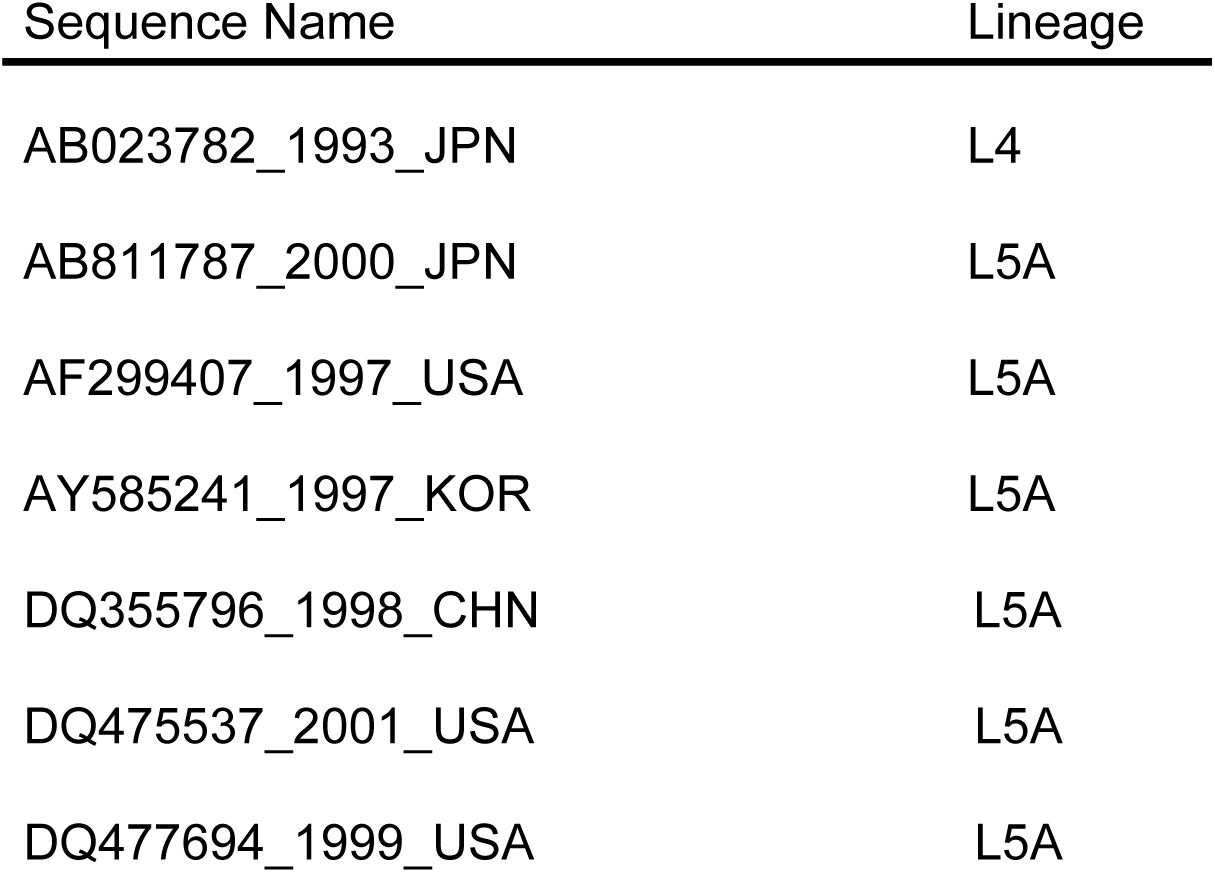
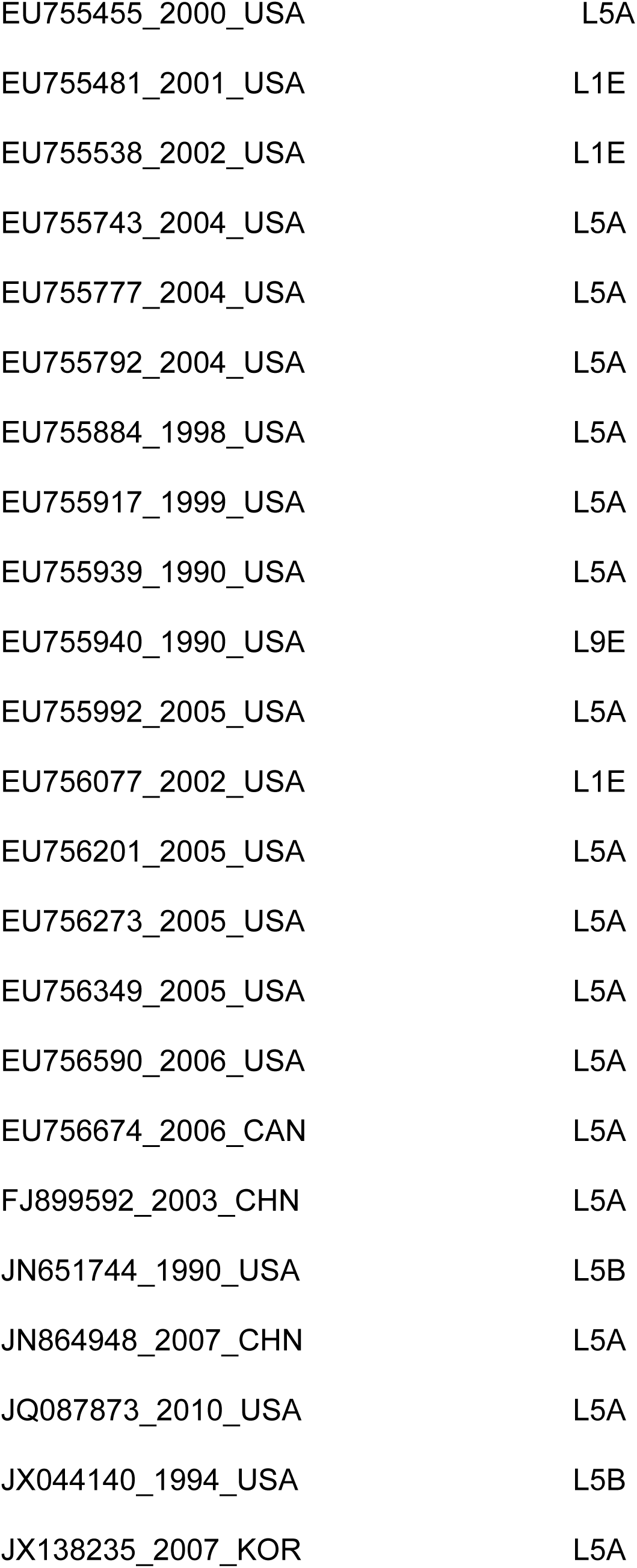
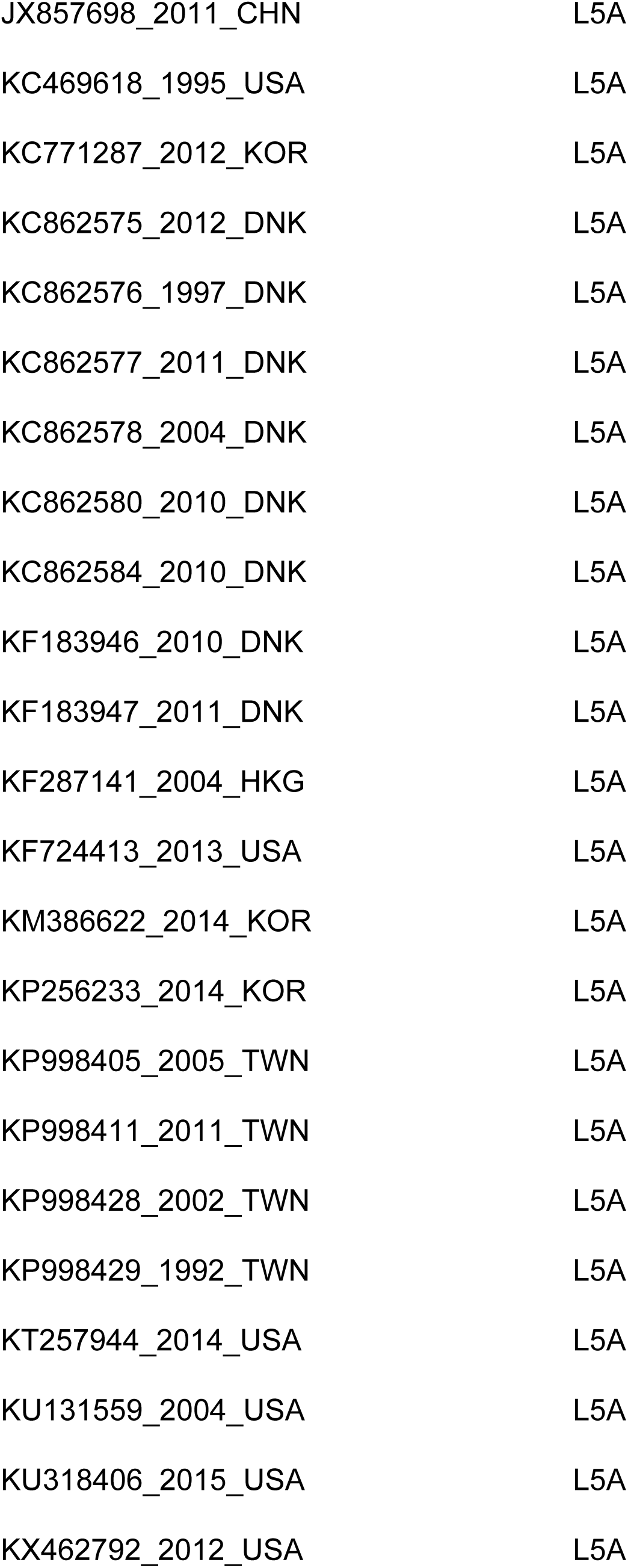
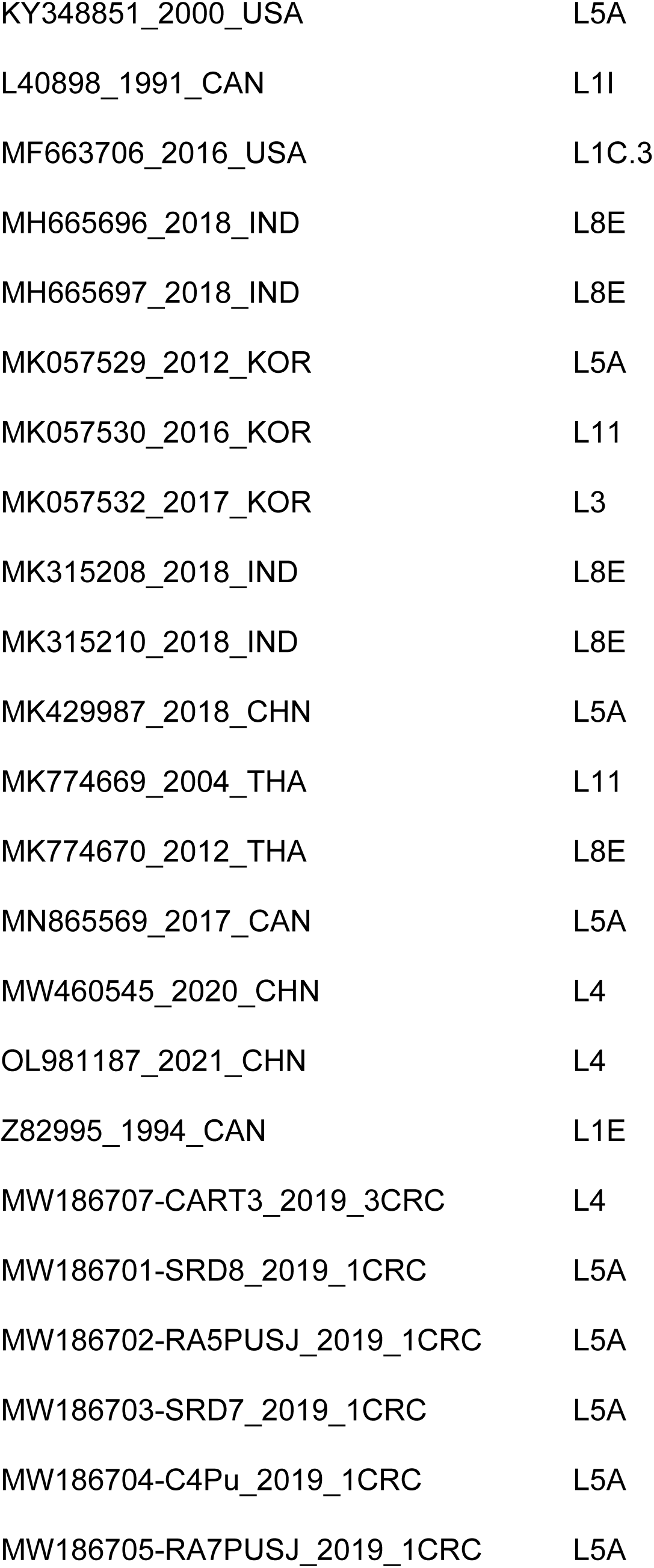
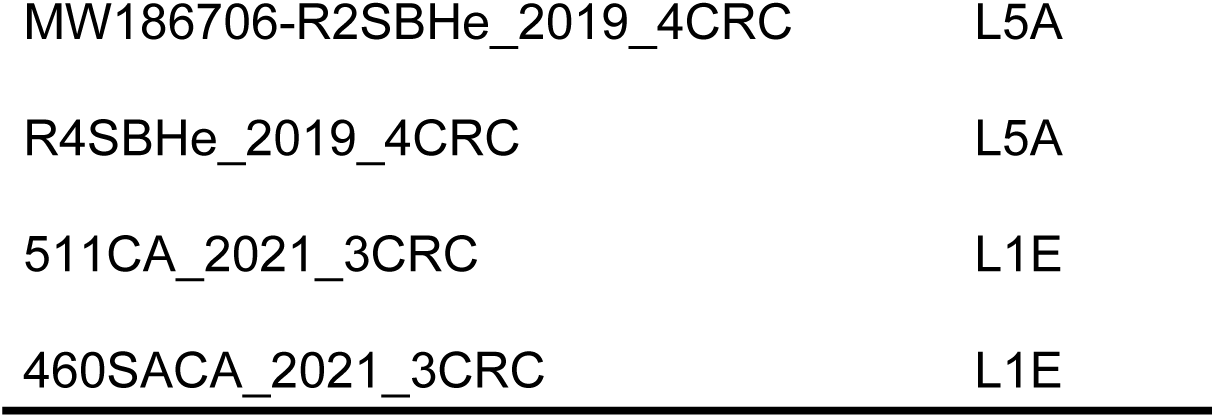
Lineage classification of the analyzed sequences based on the Nextclade software.

### Supplementary Figures

**Figure S1.**
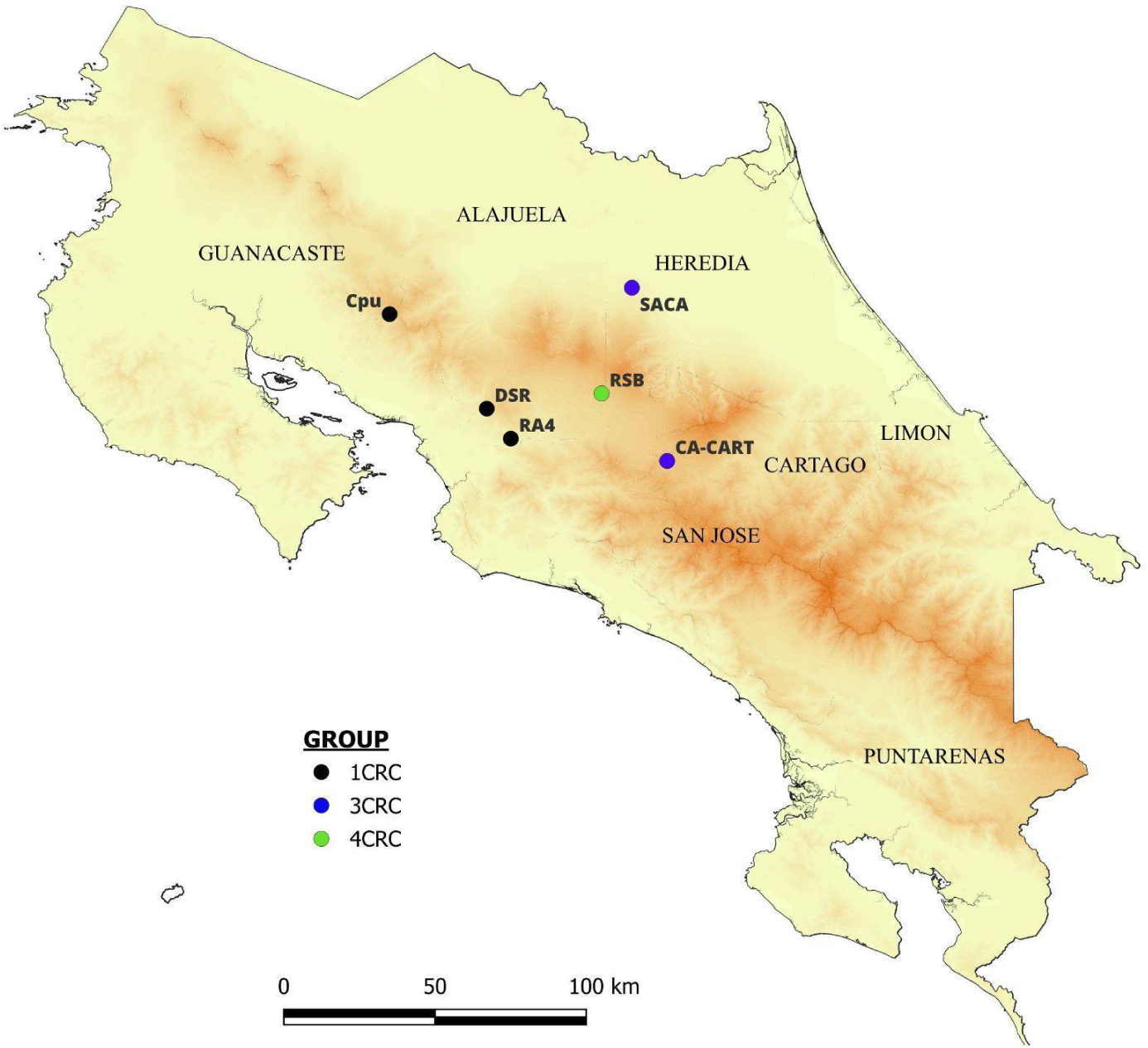
Geographical distribution of the Costa Rican PRRSV-2 clusters. The map indicates the farm locations from which samples were collected for this study. Each point is colored to correspond with the phylogenetic clusters defined in Figure 1: 1CRC (black), 3CRC (blue), and 4CRC (green). Farm codes are labeled for reference.

**Figure S2.**
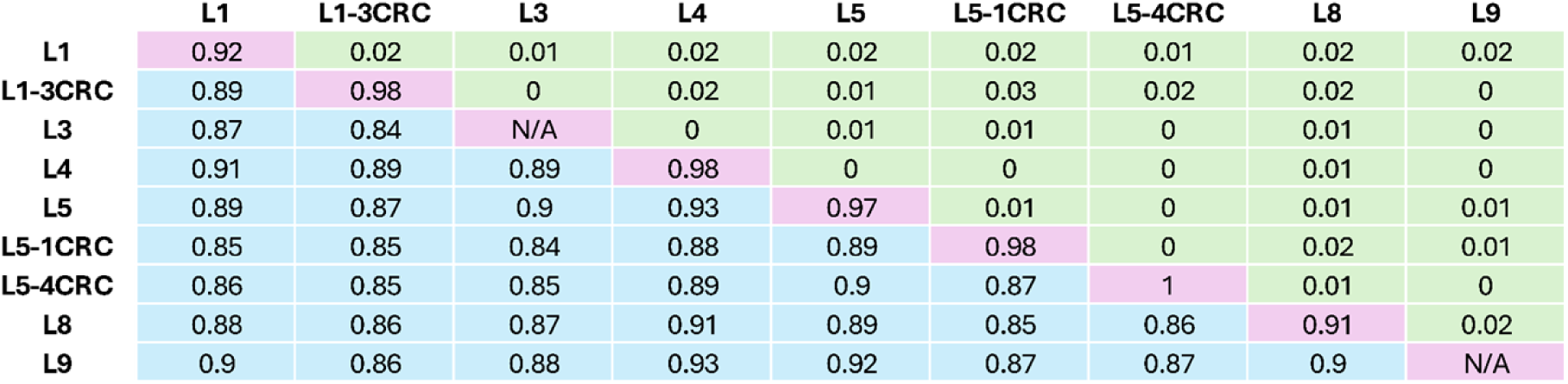
Average pairwise genetic distance among lineages. Percent genetic difference within (intra-, highlighted in magenta) and between (inter-, highlighted in light blue) sublineages L1, L3, L4, L5, L8, and L9. The standard deviation of inter-lineage distances is highlighted in green.

**Figure S3.**
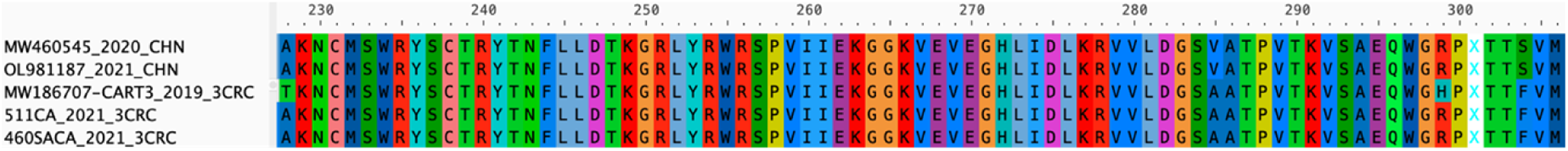
Amino acid comparison between lineages. Lineage 4 (L4) reference sequences from China and the Costa Rican sequences classified as lineage 1, sublineage E by Nextclade are shown, including the discordant sequence MW186707-CART3_2019_3CRC.

## Notes

### Competing Interest Statement

The authors have declared no competing interest.

## REFERENCES

1. Plagemann PGW. Porcine Reproductive and Respiratory Syndrome Virus : Origin Hypothesis. 2003;9: 903–908.

2. Espinoza AC, Velásquez MR. Síndrome Reproductivo y Respiratorio Porcino: Una revisión del agente etiológico y su influencia en el comportamiento actual de la enfermedad. Rev Investig Vet Perú. 2021;32: e19645. doi:10.15381/rivep.v32i1.19645

3. Amirpour Haredasht S, Polson D, Main R, Lee K, Holtkamp D, Martínez-López B. Modeling the spatio-temporal dynamics of porcine reproductive & respiratory syndrome cases at farm level using geographical distance and pig trade network matrices. BMC Vet Res. 2017;13: 163. doi:10.1186/s12917-017-1076-6

4. Shi M, Lemey P, Brar MS, Suchard MA, Murtaugh MP, Carman S, et al. The spread of type 2 porcine reproductive and respiratory syndrome virus (prrsv) in North America: A phylogeographic approach. Virology. 2013;447: 146–154. doi:10.1016/j.virol.2013.08.028

5. Nelson MI, Viboud C, Vincent AL, Culhane MR, Detmer SE, Wentworth DE, et al. Global migration of influenza A viruses in swine. Nat Commun. 2015;6: 6696. doi:10.1038/ncomms7696

6. McCullough S, Gorcyca D, Chladek D. Isolation of swine infertility and respiratory syndrome virus (isolate ATCC VR-2332) in North America and experimental reproduction of the disease in gnotobiotic pigs. J Vet Diagn Invest. 1992;4: 117–126. doi:10.1177/104063879200400201

7. Wensvoort G, Terpstra C, Pol JMA, Laak EA, Bloemraad M, Kluyver EP de, et al. Mystery swine disease in the Netherlands : The isolation of Lelystad virus. Vet Q. 1991;13 (3): 121–130. doi:10.1080/01652176.1991.9694296

8. Adams MJ, Lefkowitz EJ, King AMQ, Harrach B, Harrison RL, Knowles NJ, et al. Changes to taxonomy and the International Code of Virus Classification and Nomenclature ratified by the International Committee on Taxonomy of Viruses (2017). Arch Virol. 2017;162: 2505–2538. doi:10.1007/s00705-017-3358-5

9. Kappes MA, Faaberg KS. PRRSV structure, replication and recombination: Origin of phenotype and genotype diversity. Virology. 2015;479–480: 475–486. doi:10.1016/j.virol.2015.02.012

10. Kimman TG, Cornelissen LA, Moormann RJ, Rebel JMJ, Stockhofe-Zurwieden N. Challenges for porcine reproductive and respiratory syndrome virus (PRRSV) vaccinology. Vaccine. 2009;27: 3704–3718. doi:10.1016/j.vaccine.2009.04.022

11. Popescu LN, Trible BR, Chen N, Rowland RRR. GP5 of porcine reproductive and respiratory syndrome virus (PRRSV) as a target for homologous and broadly neutralizing antibodies. Vet Microbiol. 2017;209: 90–96. doi:10.1016/j.vetmic.2017.04.016

12. Brar MS, Shi M, Ge L, Carman S, Murtaugh MP, Leung FC-C. Porcine reproductive and respiratory syndrome virus in Ontario, Canada 1999 to 2010: genetic diversity and restriction fragment length polymorphisms. J Gen Virol. 2011;92: 1391–1397. doi:10.1099/vir.0.030155-0

13. Guzman Saborío M. Prevalencia y caracterización molecular del Virus del Síndrome Reproductivo y Respiratorio Porcino (PRRSV) en cerdos de producción de Costa Rica. Nacional. 2020.

14. Meléndez R, Guzmán M, Jiménez C, Piche M, Jiménez E, León B, et al. Seroprevalence of porcine reproductive and respiratory syndrome virus on swine farms in a tropical country of the Middle Americas: the case of Costa Rica. Trop Anim Health Prod. 2021;53: 441. doi:10.1007/s11250-021-02799-9

15. Guzmán M, Meléndez R, Jiménez C, Piche M, Jiménez E, León B, et al. Analysis of ORF5 sequences of Porcine Reproductive and Respiratory Syndrome virus (PRRSV) circulating within swine farms in Costa Rica. BMC Vet Res. 2021;17: 1–11. doi:10.1186/s12917-021-02925-7

16. Altschul SF, Gish W, Miller W, Myers EW, Lipman DJ. Basic local alignment search tool. J Mol Biol. 1990;215: 403–10. doi:10.1016/S0022-2836(05)80360-2

17. Hall TA. BioEdit: a user-friendly biological sequence alignment editor and analysis program for Windows 95/98/NT. Nucl Acids Symp. 1999;41: 95–98.

18. Nguyen LT, Schmidt HA, Haeseler AV, Minh BQ. IQ-TREE: A fast and effective stochastic algorithm for estimating maximum-likelihood phylogenies. Mol Biol Evol. 2015;32: 268–274. doi:10.1093/molbev/msu300

19. Rambaut A, Lam TT, Carvalho LM, Pybus OG. Exploring the temporal structure of heterochronous sequences using TempEst (formerly Path-O-Gen). Virus Evol. 2016;2: vew007. doi:10.1093/ve/vew007

20. Martin DP, Murrell B, Golden M, Khoosal A, Muhire B. RDP4: Detection and analysis of recombination patterns in virus genomes. Virus Evol. 2015;1: 1–5. doi:10.1093/ve/vev003

21. Drummond AJ, Rambaut A. Bayesian Evolutionary Analysis Sampling Trees v1.0. 2003; 79–96.

22. Drummond AJ, Ho SYW, Phillips MJ, Rambaut A. Relaxed phylogenetics and dating with confidence. PLoS Biol. 2006;4: 699–710. doi:10.1371/journal.pbio.0040088

23. Suchard MA, Lemey P, Baele G, Ayres DL, Drummond AJ, Rambaut A. Bayesian phylogenetic and phylodynamic data integration using BEAST 1.10. Virus Evol. 2018;4: 1–5. doi:10.1093/ve/vey016

24. Kahle D, Wickham H. ggmap: Spatial Visualization with ggplot2. R J. 2013. Available: https://digitalcommons.unl.edu/r-journal/397

25. Aksamentov I, Roemer C, Hodcroft EB, Neher RA. Nextclade: clade assignment, mutation calling and quality control for viral genomes. J Open Source Softw. 2021;6: 3773. doi:10.21105/joss.03773

26. Yim-im W, Anderson TK, Paploski IAD, VanderWaal K, Gauger P, Krueger K, et al. Refining PRRSV-2 genetic classification based on global ORF5 sequences and investigation of their geographic distributions and temporal changes. Microbiol Spectr. 2023;11: e02916–23. doi:10.1128/spectrum.02916-23

27. Muhire BM, Varsani A, Martin DP. SDT: A Virus Classification Tool Based on Pairwise Sequence Alignment and Identity Calculation. Kuhn JH, editor. PLoS ONE. 2014;9: e108277. doi:10.1371/journal.pone.0108277

28. Shi M, Lam TTY, Hon CC, Hui RKH, Faaberg KS, Wennblom T, et al. Molecular epidemiology of PRRSV: A phylogenetic perspective. Virus Res. 2010;154: 7–17. doi:10.1016/j.virusres.2010.08.014

29. Holck J, Polson D. The Financial Impact of PRRS Virus. 2003.

30. Cho JG, Dee SA. Porcine reproductive and respiratory syndrome virus. Theriogenology. 2006;66: 655–662. doi:10.1016/j.theriogenology.2006.04.024

31. Corzo CA, Mondaca E, Wayne S, Torremorell M, Dee S, Davies P, et al. Control and elimination of porcine reproductive and respiratory syndrome virus. Virus Res. 2010;154: 185–192. doi:10.1016/j.virusres.2010.08.016

32. Pileri E, Mateu E. Review on the transmission porcine reproductive and respiratory syndrome virus between pigs and farms and impact on vaccination. Vet Res. 2016;47: 108. doi:10.1186/s13567-016-0391-4

33. Thomann B, Rushton J, Schuepbach-Regula G, Nathues H. Modeling Economic Effects of Vaccination Against Porcine Reproductive and Respiratory Syndrome: Impact of Vaccination Effectiveness, Vaccine Price, and Vaccination Coverage. Front Vet Sci. 2020;7. doi:10.3389/fvets.2020.00500

34. Alkhamis MA, Perez AM, Murtaugh MP, Wang X, Morrison RB. Applications of Bayesian phylodynamic methods in a recent U.S. porcine reproductive and respiratory syndrome virus outbreak. Front Microbiol. 2016;7: 1–10. doi:10.3389/fmicb.2016.00067

35. Kapur V, Elam MR, Pawlovich TM, Murtaugh MP. Genetic variation in porcine reproductive and respiratory syndrome virus isolates in the midwestern United States. J Gen Virol. 1996;77 ( Pt 6): 1271–1276. doi:10.1099/0022-1317-77-6-1271

36. Arruda AG, Tousignant S, Sanhueza J, Vilalta C, Poljak Z, Torremorell M, et al. Aerosol Detection and Transmission of Porcine Reproductive and Respiratory Syndrome Virus (PRRSV): What Is the Evidence, and What Are the Knowledge Gaps? Viruses. 2019;11: 712. doi:10.3390/v11080712

37. Havas KA, Brands L, Cochrane R, Spronk GD, Nerem J, Dee SA. An assessment of enhanced biosecurity interventions and their impact on porcine reproductive and respiratory syndrome virus outbreaks within a managed group of farrow-to-wean farms, 2020–2021. Front Vet Sci. 2023;9. doi:10.3389/fvets.2022.952383

38. Nan Y, Wu C, Gu G, Sun W, Zhang Y-J, Zhou E-M. Improved Vaccine against PRRSV: Current Progress and Future Perspective. Front Microbiol. 2017;8: 1635. doi:10.3389/fmicb.2017.01635

39. Mondaca E, Batista L, Cano J-P, Díaz E, Philips R, Polson D. General guidelines for porcine reproductive and respiratory syndrome regional control and elimination projects. J Swine Health Prod. 2014;22: 84–88.

40. Lo Presti A, Cella E, Giovanetti M, Lai A, Angeletti S, Zehender G, et al. Origin and evolution of Nipah virus. J Med Virol. 2016;88: 380–388. doi:10.1002/jmv.24345

